# Blue light is a universal tactic signal for *Escherichia coli*

**DOI:** 10.1101/211474

**Authors:** Tatyana Perlova, Martin Gruebele, Yann R. Chemla

**Affiliations:** Department of Physics, University of Illinois at Urbana-Champaign; Department of Chemistry, University of Illinois at Urbana-Champaign; Center for the Physics of Living Cells, University of Illinois at Urbana-Champaign

## Abstract

Blue light has been shown to elicit a tumbling response in *E. coli*, a non-phototrophic bacterium. The exact mechanism of this phototactic response is still unknown, and its biological significance remains unclear. Here, we quantify phototaxis in *E. coli* by analyzing single-cell trajectories in populations of free-swimming bacteria before and after light exposure. Bacterial strains expressing only one type of chemoreceptor reveal that all five *E. coli* receptors - Aer, Tar, Tsr, Tap and Trg - are capable of mediating a response to light. In particular, light exposure elicits a running response in Tap-only strain, the opposite of the tumbling response observed for all other strains. Light therefore emerges as a universal stimulus for all *E. coli* chemoreceptors. We also show that blue light exposure causes a reversible decrease in swimming velocity, a proxy for proton motive force. We hypothesize that rather than sensing light directly, chemoreceptors sense light-induced perturbations in proton motive force.

**Importance:** Our findings provide new insights on the mechanism of *E. coli* phototaxis, showing that all five chemoreceptor types respond to light and that their interactions play an important role in cell behavior. Our results also open up new avenues for examining and manipulating *E. coli* taxis. Since light is a universal stimulus, it may provide a way to quantify interactions between different types of receptors. Since light is easier to control spatially and temporally than chemicals, it may be used to study swimming behavior in complex environments. Since phototaxis can cause migration of *E. coli* bacteria in light gradients, light may be used to control bacterial density for studying density-dependent processes in bacteria.

## Introduction

Phototaxis, the light-dependent movement of microorganisms, was first reported as early as the 19th century in certain species of purple bacteria (1). Along with halobacteria and cyanobacteria, purple bacteria are phototrophic, i.e. they use light as a source of energy. Phototaxis confers an obvious advantage to phototrophic bacteria as it allows them to migrate to optimal illumination conditions (2, 3). *Escherichia coli*, on the other hand, is a surprising example of a non-phototrophic bacterium for which exposure to blue light results in changes in motile behavior (4–7).

*E. coli* motility is governed by a few simple principles that allow it to find the most favorable environment efficiently. *E. coli* is propelled by a bundle of helical, rotating flagella and swims by alternating between two types of motion—‘runs’, during which cells swim in one direction along an approximately straight path, and ‘tumbles’, during which cells randomly reorient. Runs correspond to counterclockwise (CCW) rotation of all flagellar motors, which results in a tight bundle of flagella propelling the cell forward. During tumbles, one or more flagella rotate clockwise (CW), breaking from the bundle and causing random reorientation of the cell before the next run (8). The fraction of time spent tumbling—the ‘tumble bias’—therefore depends on the fraction of time each flagellar motor rotates CW. Tumble bias changes in response to intracellular cues such as proton motive force (PMF) (9), or extracellular chemical cues (10), resulting in a behavior known as ‘taxis’.

This behavior is controlled by a simple signaling network (Fig. 1A) (10). The intracellular signaling molecule CheY in its active, phosphorylated form (CheY-P), binds to the flagellar motors, causing a switch in rotational direction from CCW to CW. CheY is phosphorylated by the kinase CheA and dephosphorylated by the phosphatase CheZ. *E. coli* has five types of transmembrane receptors (Fig. 1A) of varying abundance—Tar, Tsr, Aer, Tap, and Trg—that sense a range of environmental signals (11, 12). These receptors form complexes with CheA, coupling its kinase activity to environmental conditions. For example, binding of chemical repellents to the receptors’ ligand-binding domains results in increased CheA activity, a higher concentration of CheY-P, a higher probability of CW motor rotation, and therefore a higher tumble bias. *Vice versa*, binding of chemoattractants deactivates CheA, resulting in lower tumble bias. Finally, CheA activity is also regulated by receptor methylation, which is controlled by methyltransferase CheR and methylesterase CheB, the latter of which is active only when phosphorylated by active CheA (10) (Fig. 1A). CheR and CheB regulation of CheA activity formsa negative feedback loop that allows bacteria to adapt to new conditions. The net result is that *E. coli* run lengths increase as conditions become more favorable, cells migrate towards a better environment, and eventually adapt as the tumble bias returns to its basal value (13).

**Figure 1.**
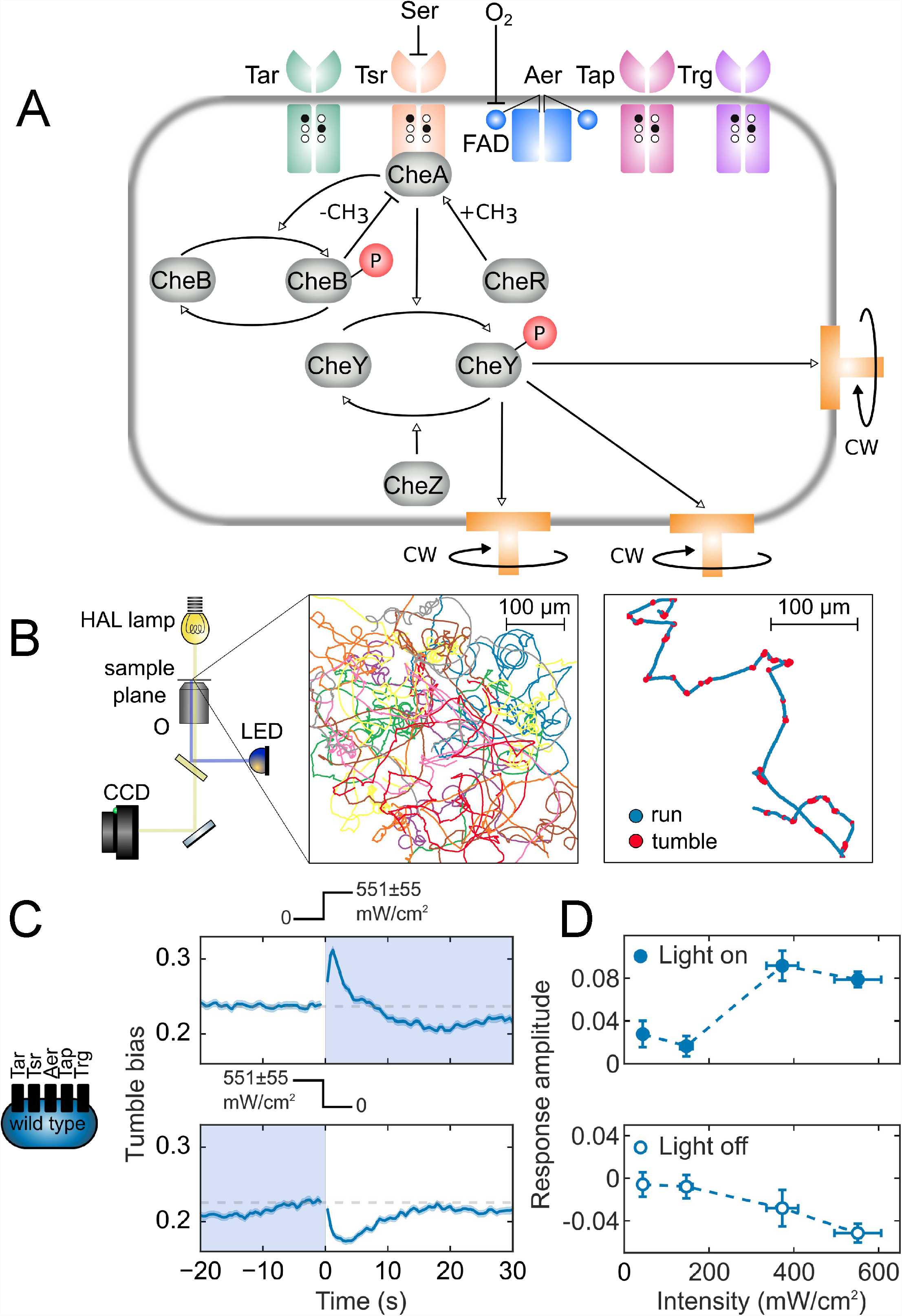
Phototaxis in *E. coli.* (A) Schematic of the chemotaxis network. Five types of chemoreceptors (Tar, Tsr, Aer, Trg, and Tap) sensitive to different extra- and intracellular cues modulate the activity of the kinase CheA, which phosphorylates the signaling molecule CheY. Phosphorylated CheY binds to flagellar motors causing them to switch to clockwise (CW) rotation. Chemotactic adaptation is mediated by methyltransferase CheR and methylesterase CheB. Methylation of receptor sites (black circles) by CheR increases kinase activity. CheB, when phosphorylated by active CheA, de-methylates receptors (white circles) and decreases kinase activity. (B) Experimental and data analysis framework for studying phototaxis in *E. coli*. An inverted light microscope with 20x objective (O) with wide-range HAL lamp (yellow light path) images swimming cells, and a blue LED (blue light path) stimulates cells. Movies of swimming bacteria are captured by a CCD camera. Bacteria are detected in each movie frame and their coordinates are linked into trajectories, which are filtered and analyzed to assign runs (blue lines) and tumbles (red circles). (C) Response of the wild-type strain RP437 (schematic indicates that all receptor types are present) to a turn-on and turn-off in blue light of intensity 551 ± 55 mW/cm^2^. Light exposure is indicated by the shaded area as well as by the light intensity profiles above the plots. Each point on the time trace is the average of the tumble biases of ∼6000-7000 trajectories. The shading around the time trace represents the standard error of the mean tumble bias. (D) Amplitude of the response to turn-on (filled circles) and turn-off (empty circles) as a function of light intensity.

The taxis signaling network is characterized by its (1) extreme sensitivity—an *E. coli* cell can respond to concentration changes as small as ∼3 nM, corresponding to just a few molecules per cell volume (14); (2) wide dynamic range—a cell is sensitive to changes of up to 5 orders of magnitude in concentration (10); and (3) the ability to integrate diverse extracellular cues—not just concentrations of various chemicals (’chemotaxis’), but temperature (’thermotaxis’), pH (’pH-taxis’), and light (’phototaxis’) (12, 15, 16). Chemotaxis in *E. coli* has been studied extensively and serves as a paradigm for the way living cells modulate their behavior in response to environmental signals (13, 17–20). However, there have been only a handful of studies on *E. coli* phototactic response (4–7), and the adaptive value of phototaxis remains unclear.

Here, we study phototaxis by analyzing single-cell trajectories in populations of *E. coli* bacteria free-swimming in 2D, before and after exposure to blue light. Our results show that light is a universal tactic signal and elicits responses mediated by all 5 types of receptors. Single-receptor mutant measurements confirm that Tar and Aer receptors mediate increased tumbling in bacteria exposed to light, in agreement with prior studies (7). The role of the other three receptors in phototaxis was previously unknown. We find that Tsr and Trg also mediate tumbling in response to light, whereas Tap mediates a running response. Despite Tap being a low abundance receptor, we observe that several multi-receptor strains containing Tap exhibit running responses to light. A reversible decrease in bacterial swimming velocity that we observe upon light exposure suggests that light perturbs electron transport and/or proton motive force (PMF). Based on these results, we propose a mechanism for a universal tactic response to light in which *E. coli* receptors sense light-induced perturbations of PMF or parameters coupled to PMF.

## Results

### Response to blue light requires at least one receptor, functional CheY, CheR and CheB

We analyzed swimming trajectories from thousands of cells stimulated by a turn-on or turn-off of blue light (Fig. 1B; Supplementary Fig. 1-3; Materials and Methods). Similarly to Wright *et al.* (7), we observed wild-type *E. coli* tumble more in response to a turn-on, then return to the prestimulus tumble bias, and exhibit the opposite response to a turn-off (Fig. 1C). The light intensities needed to elicit a detectable response were similar to those reported by Taylor and Koshland (4) and by Taylor *et al.* (5), but were significantly larger than those of Wright and co-workers—more than 300 mW/cm^2^ compared to 7 mW/cm^2^ (7). We attribute the discrepancy to different growth conditions in the latter study (Materials and Methods). When using the same substrates in growth and motility media as Wright *et al.* (M9 supplemented with 5 mg/ml glycerol and motility buffer with 5 mM lactate) we reproduced similar responses at a lower light intensity (Supplementary Fig. 4). Under our conditions (M9 supplemented with 4 mg/ml succinate and motility buffer with 4 mg/ml succinate), the response amplitude increased with light intensity and saturated at ∼400 mW/cm^2^ (Fig. 1C).

We confirmed that this response is mediated by the chemotactic network by performing control experiments with mutants missing different components of the network (Supplementary Fig. 5). We observed no response to light in either a receptorless strain (UU1250, Table 1) or a strain lacking functional CheY (CR20, Table 1). Therefore, similar to other types of tactic stimuli, blue light modulates the activity of receptor signaling complexes, and this signal is communicated to the flagellar motors through the signaling molecule CheY-P. Blue light does not appear to affect switching of flagellar motor rotation directly.

**Table 1.** Strains used in this work

| Strain | Genotype | Comments | Source |
| --- | --- | --- | --- |
| RP437 |  | Wild-type chemotaxis strain | (50) |
| $\Delta R$ | $\Delta cheR$ | | Christopher Rao |
| $\Delta B$ | $\Delta cheB$ | | Christopher Rao |
| CR20 | $CheY::FRT$ | Runner | Christopher Rao (51) |
| UU1250 | $\Delta(tar-tap), \Delta tsr, \Delta aer, \Delta trg$ | Receptorless strain | John S. Parkinson (26) |
| UU1615 | $\Delta aer \Delta tar \Delta tap \Delta trg$ | Tsr-only strain | John S. Parkinson (52) |
| UU1624 | $\Delta aer \Delta tsr \Delta tap \Delta trg$ | Tar-only strain | John S. Parkinson (52) |
| UU1250 + pSB20 | $\Delta(tar-tap) \Delta tsr \Delta aer, \Delta trg$ | Aer-only strain, contains pSB20 plasmid | John S. Parkinson (7) |
| UU1250 + pPA705 | $\Delta(tar-tap) \Delta tsr \Delta aer, \Delta trg$ | Trg-only strain, contains pPA705 plasmid | This work |
| TP6 | $\Delta(tar-tap) \Delta tsr \Delta aer \Delta trg$ | Tap-only strain, contains pTP1 plasmid | This work |
| RP8604 | $\Delta tsr \Delta trg$ | Tar-Aer-Tap strain | John S. Parkinson |
| RP1131 | $trg::Tn10$ | $\Delta Trg$ strain | John S. Parkinson |
| UU1117 | $\Delta aer$ | $\Delta Aer$ strain | John S. Parkinson (53) |

We also confirmed that the observed adaptation to blue light is mediated by chemotactic network proteins CheR and CheB. A strain lacking the receptor demethylase CheB (ΔB, Table 1) had a very high tumble bias which did not increase measurably upon light exposure (Supplementary Fig. 5). The lack of response indicates that fully methylated receptors cannot be further activated by light. A strain lacking the receptor methyltransferase CheR (ΔR, Table 1) exhibited an initial sharp increase in tumble bias, similarly to the wild-type strain, but the tumble bias then failed to return to a lower value (Supplementary Fig. 5). The lack of adaptation is expected: in the absence of CheR, receptor methylation level is low and adaptation through de-methylation is therefore impossible. In summary, the lack of response in the ΔCheB strain indicates that a tumbling response to light is caused by receptor activation in the wild-type strain. The post-response kinetics in the ΔCheR strain indicate that adaptation in the wild-type strain is mediated by CheR through the negative feedback loop of the chemotaxis network. (Note that rather than simply leveling off, the tumble bias of ΔCheR continues to increase slowly, which we speculate is due to a slow, secondary effect.)

### All five *E. coli* chemoreceptors mediate blue light-induced changes in tumble bias

#### Turn-on responses

The results above demonstrate that the blue light response in wild-type *E. coli* requires functioning receptor complexes. To determine the independent contribution of individual receptor types, we measured the responses of mutants expressing only a single receptor type (Materials and Methods). Abundances (i.e. copy numbers) of different receptor types can span two orders of magnitude in wild-type strains (21). Therefore, low-copy-number receptors Aer, Tap, and Trg were expressed from a plasmid, with the concentration of inducer adjusted to ensure that a tumble bias in the mutant strain in the absence of stimulus was similar to that of the wild-type strain. Confirming the results of Wright *et al.* (7), we observed tumbling responses in Tar- and Aer-only strains (Fig. 2A). We also detected responses in Tsr-, Trg-, and Tap-only strains. The Tsr-only strain exhibited a weak but consistent tumbling response, and the Trg-only strain showed a tumbling response with delayed onset (Fig. 2A).

The Tap-only strain showed a unique response pattern compared to the other receptors. Tap, like Trg, can be methylated, but lacks the NWETF motif that recruits the methyltransferase CheR (22). Therefore, we expected Tap to have a low methylation level, and hence low activity in the absence of other receptors. Indeed, we observed that the Tap-only strain had a significantly lower tumble bias than the wild-type strain and did not respond to blue light exposure. The lack of response suggests that unlike Trg, Tap receptor is not activated by light exposure. In order to explore the possibility that Tap can be deactivated by light exposure and therefore mediate a running response, we performed light response measurements of the Tap-only mutant in the presence of phenol. Phenol is a repellent for Tap and therefore activates Tap receptors, increasing its tumble bias (23). Indeed, the Tap-only strain had a higher tumble bias in the presence of phenol which increased with phenol concentration. We observed running responses to light in both 2.5 mM (Fig. 2A and Supplementary Fig. 6) and 5 mM phenol (Supplementary Fig. 2). Phenol does not absorb in the blue region of the spectrum (24), which rules out the possibility that Tap responds to excited-state phenol rather than to light itself. In further controls with the Tar-only strain in the presence of 2.5 mM phenol (Supplementary Fig. 6), we observed qualitatively the same response to light turn-on and turn-off as in the absence of phenol, although the time scales of the response and adaptation were affected. These results taken together indicate that unlike the Tar, Tsr, Aer, and Trg receptors, the Tap receptor is deactivated by light exposure and mediates a running response in *E. coli*.

**Figure 2.**
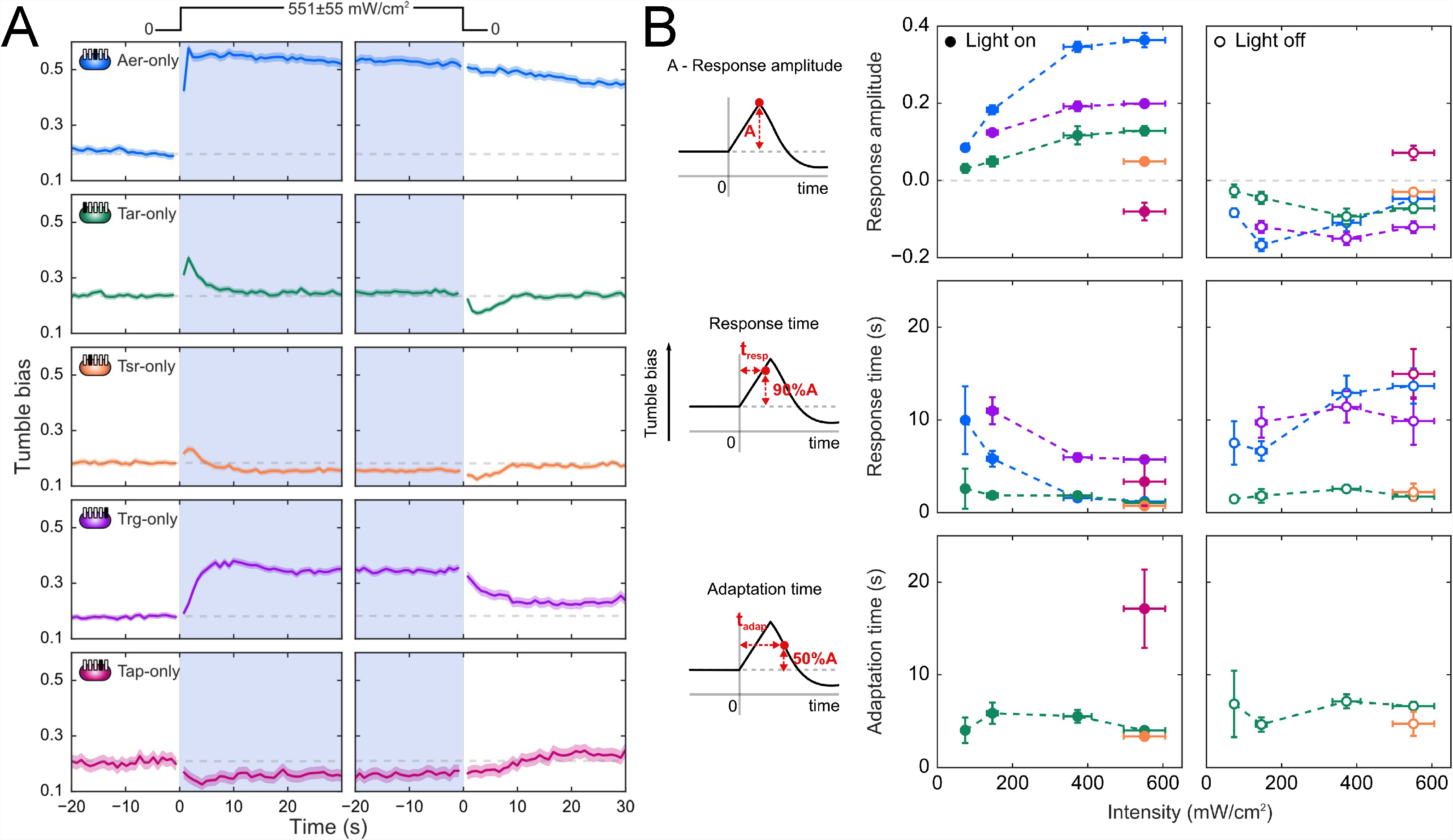
Blue light response for single-receptor *E. coli* strains. (A) Tumble bias traces for single-receptor *E. coli* strains (Table 1): Aer-only (blue), Tar-only (green), Tsr-only (orange), Trg-only (purple) and Tap-only (magenta). ∼200 - 2000 trajectories were used to calculated average tumble bias at each time point. Schematics indicate which receptor is present (filled rectangles) and absent (empty rectangles) in each strain. All responses were measured at a blue light intensity of 551 ± 55 mW/cm^2^ as indicated by the light intensity profile. Grey dashed lines show the prestimulus tumble bias for turn-on and turn-off responses. (B) Blue light-intensity dependence of parameters for the responses to turn-on (filled circles) and turn-off (empty circles) for all strains. Response amplitude, response time, and adaptation time were calculated as illustrated by the schematics.

The observed single-receptor responses exhibited adaptation behavior consistent with what is known about the mechanisms of adaptation for different receptors (Fig. 2A). Both Tar- and Tsr-only strains adapted to a steady-state tumble bias <10 s after the light turn-on, which is consistent with methylation-dependent adaptation (Fig. 2B). In contrast, Tap- and Trg-only strains did not adapt appreciably to light turn-on, consistent with the fact that they cannot recruit CheR (22). We did observe partial, slow adaptation kinetics in both mutants, which may be due to the recently reported mechanism of motor remodeling (25) or, alternatively, may reflect internal dynamics of the processes perturbed by light. The Aer receptor adapts through an unknown methylation-independent pathway that also tends to be slower (26) and consequently, we do not observe significant adaptation for the Aer-only strain within the duration of our measurement.

#### Turn-off responses

For most of the single-receptor strains, response to light turn-off was symmetrical to the response to light turn-on (Fig. 2A). For example, Tsr-only and Tar-only strains exhibited the running responses to turn-off with the adaptation kinetics similar to turn-on responses, further confirming that adaptation is mediated by the negative feedback loop of the chemotaxis network (Fig. 2A). In Trg and Tap-only strains, responses to light turn-off were also opposite in sign to those to turn-on, but with little to no adaptation (Fig. 2A). In contrast, Aer-only strain exhibited markedly slower response kinetics when high intensity light was turned off as compared to when light was turned on (Fig. 2A). This result suggests that the effect of light goes beyond receptor activation, and that Aer may be sensing a secondary effect from which it takes some time to recover, rather than light itself (Fig. 2A). However, at low light intensity, Aer responses to light turn-off and turn-on became symmetric (Supplementary Fig. 7). This intensity dependence suggests that Aer has a sensing mechanism qualitatively different at low and high light intensities.

#### Response time analysis

We also analyzed the response times of the various strains— defined as the time from the application of the stimulus to the response peak. We switched light on and off in less than one movie frame (0.08 s), and conformational changes of receptors in response to stimuli are known to happen on a sub-second time scale (27). Response to light exposure was immediate within the time resolution of the experiment for Tar, Tsr and Aer-only strains at high light intensity, while both Tap and Trg-only strains demonstrated gradual responses (Fig. 2A). Slower response times may indicate that receptors are responding to the light-induced perturbations of cellular processes on longer time scales, rather than responding directly to light. Our measurements cannot distinguish whether the observed gradual response kinetics result from a uniformly slower response across the cell population or from immediate single-cell level responses with varying time delays (28). Single-cell taxis assays that provide long (∼100 s) tumble bias traces for individual bacteria may allow distinguishing between these two explanations (28).

Figure 2B summarizes the dependence of the response amplitudes, response times, and adaptation times on light intensity for representative strains. There are several trends worth noting. First, the amplitude of the response to light exposure increased with intensity and saturated at around 400 mW/cm^2^, similarly to wild-type (Fig. 2B, left panels). This is in contrast with the results of Wright *et al.* who reported saturation above ∼10 mW/cm^2^ (7) However, asshown above, growth conditions appear to affect the sensitivity of the response (Supplementary Fig. 4). Although the peak tumble bias at saturating light intensity for single-receptor mutants is higher than for the wild-type strain, it is still lower than that for continuously tumbling bacteria (Fig. 2A). Response times to light turn-on in Aer and Trg-only strains decreased with light intensity (Fig. 2B, middle panels). No dependence of response and adaptation times on light intensity was observed for Tar-only strain (Fig. 2B, middle and right panels).

### Contribution of individual receptors to the light response in multiple receptor strains is non-additive

Our results indicate that light is a universal tactic stimulus that affects all five *E. coli* chemotactic receptors. When all receptors respond to the same stimulus, it is far from obvious how the signals from different receptors will be integrated to produce a response in wild-type strain. Chemotaxis signaling units, formed by trimers of receptor homodimers and CheA dimers, are organized into a hexagonal lattice that serves as a structural platform for interactions between receptors (29–31). Signals from low abundance receptors are thus amplified by interactions with high abundance receptors and could still contribute to the chemotactic response (32). Therefore, the contribution of individual receptor types to the overall response could be non-additive.

As shown in Fig. 1, in a wild-type strain, in which 4 out of 5 receptors (all but Tap) mediate a tumbling response to light, the integrated result of the individual receptor contributions is a tumbling response with the amplitude comparable to that of the Tar-only strain. This suggests that, despite its low abundance, the ‘running’ receptor Tap may contribute to the response of the wild-type strain and lower the amplitude of the tumbling response.

Since Tap is the only receptor type to mediate a running response to light exposure, we asked whether it can switch the sign of the response in multi-receptor mutants, strains expressing a subset of 5 chemoreceptors (Table 1). Indeed, we observed a running response in several mutants containing the Tap receptor (Fig. 3). For example, a combination of high-abundance Tar and low-abundance Aer receptors, which both mediate tumbling responses, with low-abundance Tap receptor produced a running response as illustrated by the ΔTsrΔTrg strain (Fig. 3A). Similar to the Tap-only strain, the response in this mutant was not immediate, as it took ∼5 s to reach the minimum in tumble bias. Turning the light off also caused a tumbling response, although with much lower amplitude. We also observed a running response to a light turn-on in a ΔAer strain, with a similarly slow response time (Fig. 3A). In these strains, we expect receptor abundance to be similar to that in the wild-type strain—where the abundance of Tap is much lower than that of Tar or Tsr (Fig. 3C). Therefore, the running response must be a result of amplification of Tap-mediated signaling by other receptors.

**Figure 3.**
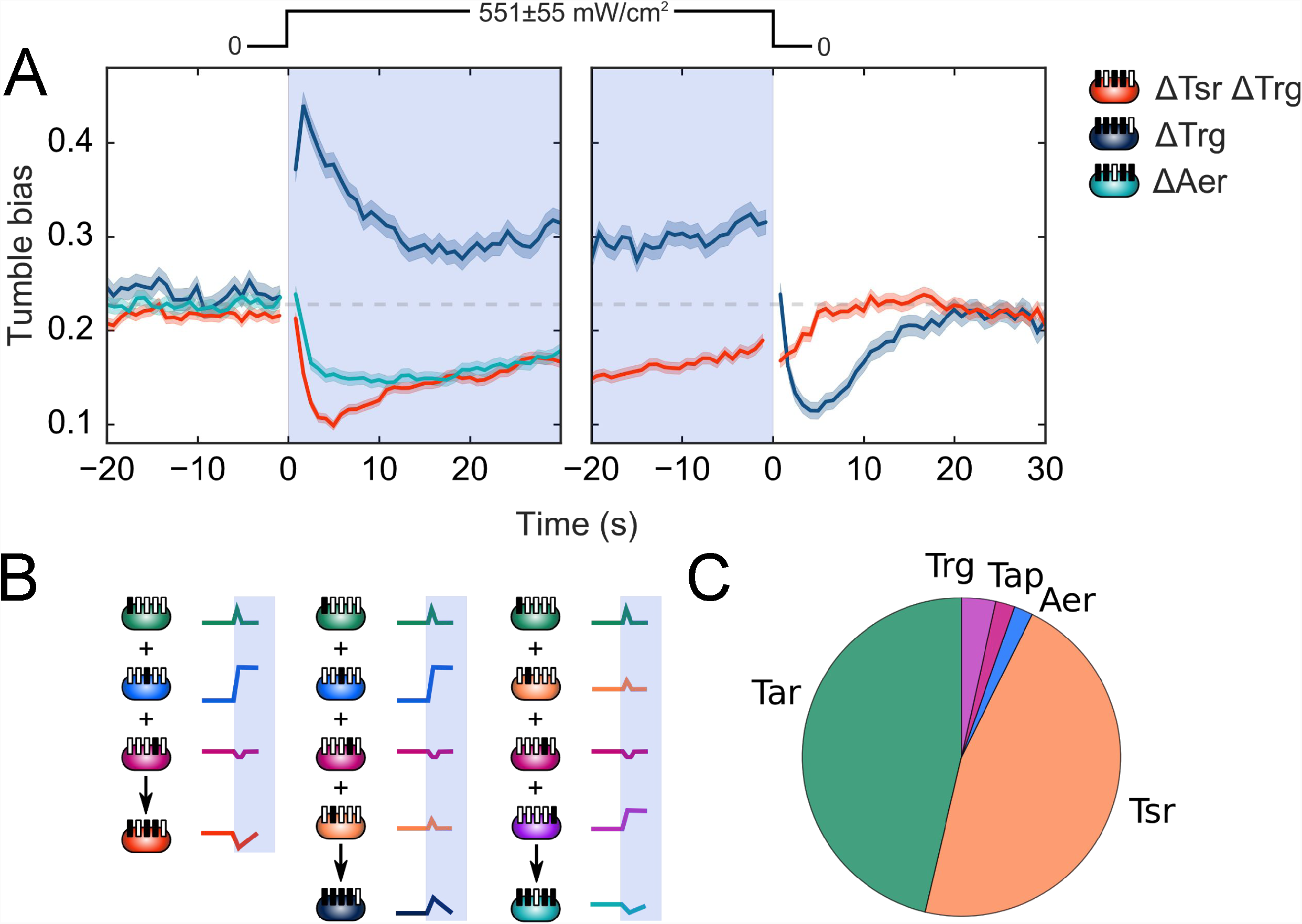
Non-additivity of individual receptor responses in blue light response of multiple-receptor strains. (A) Tumble bias traces for ΔTsrΔTrg (red), ΔTrg (dark blue) and ΔAer (cyan) strains. Schematics indicate which receptor is present in each strain. All responses were measured at a blue light intensity of 551 ± 55 mW/cm^2^ as indicated by the light intensity profile. ∼900 - 4000 trajectories were used to calculated average tumble bias at each time point. (B) Schematics of responses from single receptors that contribute to the observed responses from multiple-receptor strains. The signs and kinetics of the multiple-receptor responses are different from those expected from simple addition and is, therefore, a combined result of individual receptor responses and their interactions. (C) Pie chart of the relative abundances of different receptor types in the wild-type strain, based on data from (21).

We found that the presence of Tap was a necessary, but not sufficient condition to observe a running response. For example, adding Tsr receptor to ΔTsrΔTrg mutant resulted in a switch in response from running back to tumbling (Fig. 3A, B). This result is especially surprising given that Tsr only mediates a weak tumbling response on its own (Fig. 2). This further underlines the importance of interactions between receptors in determining the overall response.

### Blue light exposure causes a reversible decrease in swimming velocity

Despite primarily sensing different environmental signals, the 5 chemoreceptors in *E. coli* universally respond to the same blue light stimulus, one of them in the opposite direction from the other four. According to current understanding, Aer is the only receptor that binds a blue light-absorbing chromophore (flavin adenine dinucleotide, FAD) as a cofactor, and therefore could be directly photosensitive (33). Thus, the mechanism of photosensitivity is unclear for the remaining four receptors. It has been speculated that *E. coli* monitors some intracellular parameter perturbed by the absorption of blue photons. Wright and co-authors suggested, for example, that the Tar receptor may be sensing perturbations in electron transport induced by blue light, which would affect Proton Motive Force (PMF) (7). We explored whether or not the PMF hypothesis is a plausible explanation for the light sensitivity of chemoreceptors.

PMF is generated during respiration by electron and proton translocation across the membrane (34). It is used to synthesize ATP and to energize other processes in the cell such as ion transport and flagellar rotation (34). It was previously shown that the flagellar rotation rate is linearly proportional to PMF under both high and low viscous load, the latter corresponding to the load on flagella in free-swimming bacteria (35, 36). Therefore, all else being equal, the bacterial swimming velocity can serve as a proxy measure of PMF. If light does affect PMF, we can expect to see corresponding trends in swimming velocity.

Velocity traces for receptorless strain UU1250 at different light intensities are shown in Figure 4A. Light exposure caused a gradual decrease in its swimming velocity. Since the receptorless strain does not exhibit a change in tumble bias in response to light (Supplementary Fig. 4), trends in swimming velocity cannot be attributed to an imperfect run-tumble assignment (see Materials and Methods). A similar decrease in velocity was also observed for all other strains assayed, including the wild-type (Fig. 4B). The magnitude of the decrease in normalized swimming velocity after 30 s of light exposure, Δ*V*_*on*_, depends on the light intensity and varies from 0 at ≤44 mW/cm^2^ to about 8% at 550 mW/cm^2^ on average (Fig. 4B). Moreover, the velocity decrease appears reversible. About 50% of the velocity decrease is recovered within 30 s after the light is turned off at all intensity levels. It is likely that a full recovery can be achieved with sufficient waiting time, but our assay preclude measurements beyond ∼30 s. Under our experimental conditions we cannot distinguish between a velocity recovery in previously exposed bacteria and a velocity increase due to unexposed bacteria swimming in the field of view after 30 s (Materials and Methods). Nevertheless, our results strongly suggest a reversible decrease in velocity, which is consistent with reversible perturbation of PMF by blue light exposure as opposed to irreversible motor photodamage.

**Figure 4.**
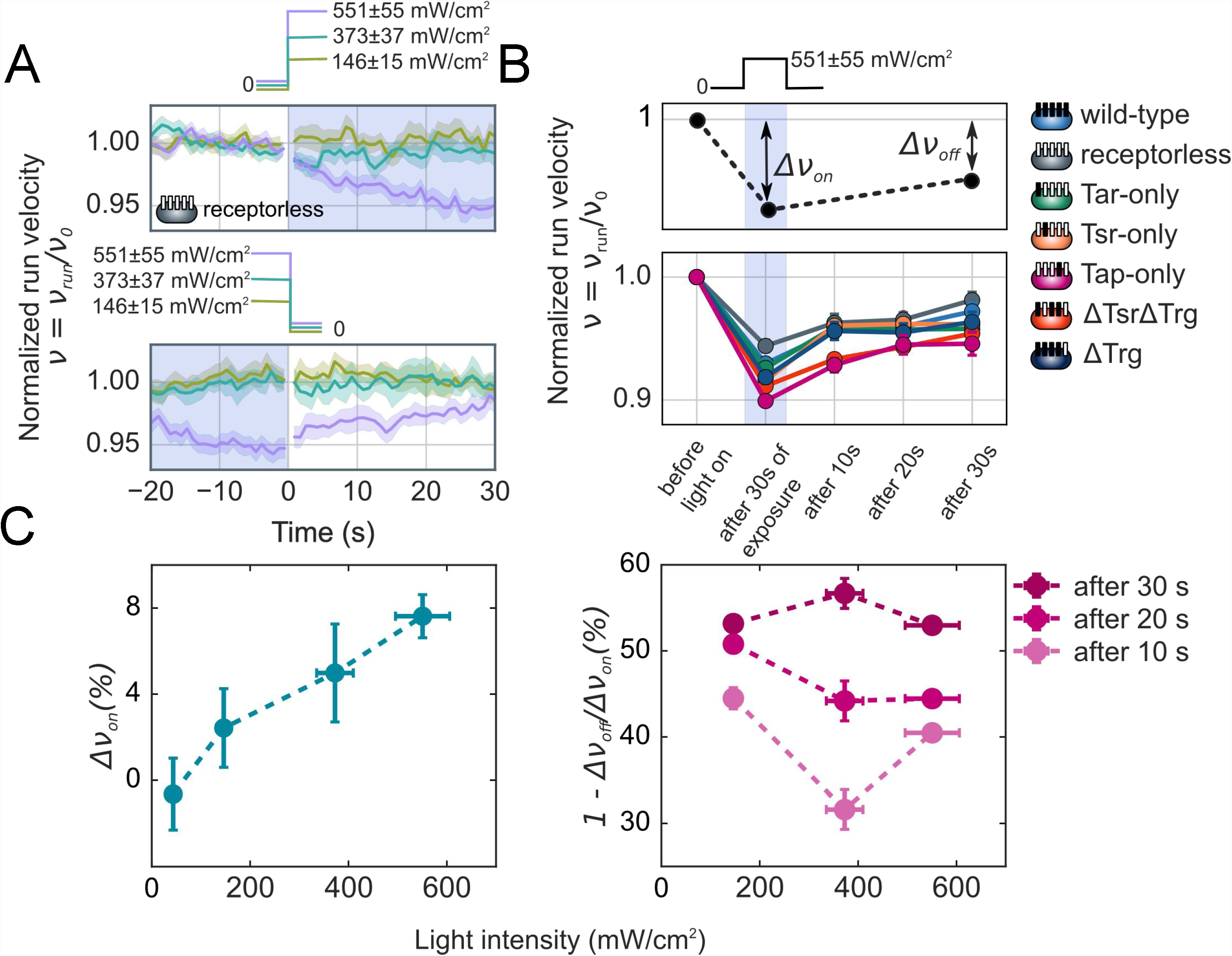
Effect of light on running velocity. (A) Light exposure causes a reversible decrease in swimming velocity. Normalized velocity traces are shown for receptorless strain UU1250. ∼2000 trajectories were used to calculated velocity at each time point. Light intensity is indicated by the trace color. The velocity is normalized by its prestimulus value (calculated in a 20 s window before light exposure; Materials and Methods). (B) Bar plot of the normalized swimming velocity for different *E. coli* strains. The normalized velocity was calculated in a 4-s window after 30 s of exposure to light, then 10, 20, and 30 s after the light was turned off. Light exposure and intensity are indicated by the intensity profile above the plot. (C) Dependence of velocity decrease and recovery on light intensity. Velocity decrease, Δ*v*_*on*_, and velocity recovery, 1 - Δ*v*_*on*_/Δ*v*_*off*_, were calculated as illustrated by the schematic in (B). Velocity recovery is shown 10, 20, and 30 s after light turn-off as indicated by the color.

## Discussion

A tumbling response to blue light in *E. coli* was first demonstrated in 1975 (4). Later, Wright *et al.* (7) identified two types of receptors essential for the tumbling response to blue light: Tar and Aer. Our work shows that the response to light in *E. coli* is mediated by all of the receptor types, including Tsr, Trg and Tap. Downstream from the receptors the effect of light on the chemotaxis network is consistent with that of other tactic stimuli. Light thus emerges as a universal tactic stimulus that affects all five *E. coli* chemoreceptors. While 4 out of 5 receptors mediate tumbling, or a repellent response to light, Tap receptor mediates running, or an attractant response. We find that, despite being a low-abundance receptor, Tap is capable of determining the direction of light response in multi-receptor mutants, likely aided by amplification of Tap-mediated signaling by other receptors.

Our results on the effect of light on swimming velocity are consistent with the hypothesis that blue light perturbs electron transport or affects PMF, as well as with the previous observation that phototaxis in *E. coli* requires a functioning electron transport chain (5). The mechanism of PMF perturbation is not clear and our experiments do not provide clues that would help to elucidate it. However, we speculate that light causes photoreduction of electron carriers such as FAD in the cytoplasmic pool, thereby disrupting electron transport. FAD is a good candidate carrier as it absorbs in the blue spectral region and is known to undergo photoconversion between its different redox and protonation states as a co-factor of LOV and BLUF flavoprotein light sensors (37–39).

The different response kinetics that we observe for single-receptor mutants may reflect different blue light sensing mechanisms of individual receptors (Figure 2 and Supplementary Fig. 7). Response times appear to fall into two categories—fast (e.g. Tar), in which the response is essentially immediate within the time resolution of our measurements and likely reflect a direct light absorption process, and slow (e.g. Trg), in which light-induced changes are presumably more indirect and take longer to have an effect. In the Aer-only strain, the response is essentially immediate at higher intensities, which is consistent with direct sensing of light by Aer through the photoreduction of its co-factor FAD (40). On the other hand, as a receptor for ‘energy-taxis,’ Aer can also sense perturbations in electron transport, proposed to occur through change in redox state of respiratory enzymes (41). We speculate that this sensing mechanism is reflected in the slower response times observed at lower light intensities (Supplementary Fig. 7). In contrast, Tsr has been shown to respond directly to changes in PMF, and may display fast response kinetics due to the low threshold needed to elicit a response (41). How the remaining receptors sense light remains unclear. Based on the response kinetics we suspect that Tar responds to light or parameters that change immediately with light intensity, while Tap and Trg sense more indirect light-induced effects (Figure 2).

To place our results in perspective, we compare the intensity values that we have used with those that bacteria may actually encounter in nature. *E. coli* bacteria live a biphasic lifecycle with their primary habitat in the mammalian gut. Between being excreted from one host and finding the next one, *E. coli* bacteria inhabit nutrient-sparse water or soil environments (42). During the environmental phase of its lifestyle, *E. coli* can be exposed to the light from the sun. The intensity of the solar illumination at the surface of Earth is ∼140 mW/cm^2^ across the visible spectrum, with roughly 60 mW/cm^2^ falling within the 60-nm band of the blue light response spectrum as measured by Wright *et al.* (7). This value is comparable to the lowest intensity at which we observed significant responses for some of the strains in this study, 74 mW/cm^2^ (Fig. 2), although not for the wild-type strain (Fig. 1). However, under different growth conditions the wild-type strain exhibits a clear response at these low light intensities (Supplementary Fig. 4) consistent with observations by Wright and co-workers (7). We speculate that the blue light response depends on growth conditions because they affect the relative abundances of different receptors types (21, 43), which influence its amplitude and sign according to our data for multi-receptor mutants (Fig. 3). Therefore, responses may be more significant for certain free-living *E. coli* strains. While it remains to be seen what the biological significance of the blue light response is, our results indicate that migration of *E. coli* bacteria due to exposure to sunlight is plausible.

We propose that blue light may be a useful tool in investigations of chemotactic behavior. Because light is a universal stimulus for all *E. coli* receptors, it may provide a way to quantify interactions between different types of receptors. In addition, light, unlike chemicals, is much easier to control both in time and in space. Therefore, using light as a stimulus may allow studying taxis behavior of bacteria in a heterogeneous environment with multiple light gradients, thereby bridging the gap between the types of gradients bacteria likely encounter in nature and those that can be generated experimentally.

## Materials and Methods

### Microbiology

#### Cell growth and media

Bacteria were grown for 20-24 hours overnight from a single colony in 1 ml of M9 minimal medium, supplemented with 4 mg/ml succinate unless otherwise noted (1x M9 salts: 12.8 g/l Na_2_HPO_4_·7H_2_O, 3g/l KH_2_PO_4_, 0.5 g/l NaCl, 1 g of NH_4_Cl; 2 μM MgSO_4_; 0.1 mM CaCl_2_; 0.5 mM of each Meth, Leu, Thr and His; 100 μg/ml thiamine; 4 mg/ml succinate) shaking at 265 RPM at 30°C with appropriate antibiotics if necessary (34 μg/l of Chloramphenicol or 100 μg/l of Ampicillin). The overnight culture was diluted 50-fold in 1 ml of the same medium and grown, shaking at 265 RPM at 30°C for 8-12 hr (to OD600 ∼0.25-0.3) with appropriate inducers if necessary. The following concentrations of inducers were used for strains with plasmids: 50 μM IPTG for UU1250 + pSB20, 0.7 μM NaSal for UU1250 + pTP1 and 0.8 μM NaSal for UU1250 + pPA705.

The over-day culture was harvested by centrifugation (1300 g, 10 min) and gently resuspended in the appropriate volume of “motility buffer” (28) (70 mM NaCl, 100 mM Tris-Cl pH 7.5, 4 mg/ml succinate, 100 μM methionine) to reach a final OD of 0.15. Bacteria were placed back in the shaker to oxygenate the media. Methionine was added to a final concentration of 100 μM prior to sample chamber assembly. Phenol was added to a final concentration of 2.5 mM or 5 mM where noted.

#### Strains

Bacterial strains and plasmids used in this study are listed in Tables 1 and 2. Plasmid pTP1 was constructed from plasmid pKG117 by subcloning the wild-type *tap* gene between NdeI-BamHI restriction sites (Table 2). Synthesis and subcloning were performed by Genscript.

**Table 2.** Plasmids used in this work

| Plasmid | Genotype | Comments | Source |
| --- | --- | --- | --- |
| pSB20 | Aer, Amp<sup>R</sup> | IPTG-inducible<br>wild-type Aer | (7) |
| pPA705 | Trg, Cm<sup>R</sup> | NaSal-inducible<br>wild-type Trg | (29) |
| pKG117 | Aer, Cm<sup>R</sup> | NaSal-inducible<br>wild-type Aer | (7) |
| pTP1 | Tap, Cm<sup>R</sup> | NaSal-inducible<br>wild-type Tap | This work |

#### Two-dimensional swimming assay

Slides (3’’ × 1’’, # 3010, Thermo-Fisher) and coverslips (22 × 22 mm, #1, VWR) were sonicated in acetone for ∼15 min, rinsed, then sonicated in KOH for 15 min, and rinsed and dried by centrifugation (1000 rpm, 3 min). Cleaning was done on the day of each experiment as we found that storing cleaned slides in distilled water even for one day resulted in the accumulation of defects on the glass surface. Prior to each experiment, slides and coverslips were passivated with bovine serum albumin (BSA, B9000S, New England BioLabs) to prevent sticking of bacteria. Slides and coverslip were incubated with 2 mg/ml BSA for ∼20 min, then rinsed with a copious amount of water and dried with nitrogen. To assemble the chamber, a drop of motility buffer (5 μl) containing *E. coli* cells was placed on a slide and gently covered with a coverslip. Care was taken to prevent the formation of air bubbles. To prevent buffer evaporation, open sides were sealed with fast curing epoxy (Devcon, 5 minute epoxy). The distance between the slide and coverslip was determined by the thickness of the liquid layer of bacterial medium and was ∼10 μm, which roughly corresponds to our 20x objective’s depth of field.

### Microscopy

We used an inverted optical microscope (Zeiss, Axio Observer A1) with a 20x objective (Zeiss, A-Plan 20x/0.45 M27) in phase contrast mode to image swimming bacteria. For observation, bacteria were illuminated from the top by a halogen lamp (HAL 12V/100W). Light from the lamp was heat filtered and passed through a 500-nm long-pass filter (Chroma, ET500lp) to exclude the possibility of excitation of bacteria by the observation light (Fig. 1b).

Excitation light from a blue LED was introduced through the back port of the microscope. A blue LED (Thorlabs, M455L3) with a collimation assembly (retaining ring, lens tube, and aspheric condenser lens (Thorlabs, ACL2520-A)) was mounted using a Zeiss Axioskop Microscope Lamphouse Port Adapter (Thorlabs, SM1A23). Excitation light passed through a 440 ± 5-nm bandpass filter (Chroma, CT440/10bp) and was directed toward the field of view by a 500-nm dichroic mirror (Chroma, 500dcxr). To achieve even illumination of the field of view, we followed the standard procedure for Koehler illumination. The microscope field stop was opened to match the field of view (≈1.2 mm in diameter for the 20x objective).

The output light intensity of the LED was determined by the current from the LED driver (Thorlabs, LEDD1B) controlled by a DAQ card (National Instruments, NI PCI-6221) and defined using a LabView interface. Neutral density filters of ND 1 and ND 0.5 were installed in the filter slider (Thorlabs, NE05B ND, and NE10B ND) to gain finer control over the resulting light intensity. Light intensity at the sample plane at different supply voltages was measured with a power meter (Newport, power meter 1916C equipped with photodiode sensor 918D-SL-OD3) placed on the microscope stage such that the illumination fitted the area of the sensor. To calculate light power density, the total power was divided by the illumination area. The unevenness of the illumination was estimated from the distribution of pixel brightness values in an image taken by the camera below its saturation and was found to be ∼10% of the mean. We used this value as an estimate for the error in light power density.

Movies of swimming bacteria were captured using a CCD camera (PointGrey, Grasshopper 3) mounted at the microscope side port. Camera calibration with USAF target 1951 (Thorlabs, R3L3S1P) gave us an estimate of the pixel size of ∼0.26 μm. The size of the area captured by the camera in the sample plane was 532 × 532 μm. Movies were recorded at a frame rate of 12 frames per second.

### Data analysis

#### Trajectory preparation and filtering

Analysis of bacterial trajectories was performed using an automated workflow implemented in Python. First, we detected bacteria in each frame of a recorded movie using the OpenCV computer vision library (44). Then, coordinates were linked into trajectories using the trackPy package (Fig. 1B) (45). At this point, we removed all trajectories shorter than 1 s, or 12 frames, from further consideration. Then we calculated instantaneous linear velocities and accelerations and angular velocities and accelerations using 1-frame windows following the procedure described by Dufour *et al.* (46).

Next, we used the following approach to filter out spurious trajectories that belonged to bacteria tethered to the surface, drifting, or swimming too slowly. For every trajectory, we calculated the average angular velocity and the 95th percentile of the linear velocity. The two-dimensional distribution of all trajectories in these coordinates contains two clusters: one cluster corresponding to normally swimming bacteria, the less populated cluster containing trajectories of very slow or surface-tethered bacteria (Supplementary Fig. 3). For each bacterial strain, we found the two coordinates of the maximum of the ‘swimming’ cluster—the most probable values of angular velocity and 95th percentile of velocity—and kept only the trajectories that lied within the a certain radius R from the maximum. With the exception of the few strains that exhibit a very strong response to light, we defined *R* as *R* = 4 <*MAD*>, where <*MAD* > is the median absolute deviation (MAD) from the maximum of the distribution, averaged across all strains with a functional chemotaxis network. Filtering was performed separately for different strains to avoid bias due to variation in the swimming behavior. We found that filtering does not affect trajectories of the bacteria that are exposed to light disproportionately: the fraction of trajectories within 4 < *MAD* > for bacteria before, during, and after light exposure is roughly the same.

The area accessible to bacteria in a slide-coverslip chamber is larger than the illuminated area. Therefore, unexposed bacteria swimming from outside of the area illuminated by blue light could subsequently swim into the field of view and affect the observed kinetics. However, the illuminated area is still larger than the observation area captured by the camera, and these bacteria have to swim 0.2 – 0.3 mm to reach the field of view. To estimate on what time scale these bacteria could contribute to the observed kinetics, we calculated the Mean Square Displacement (MSD) for trajectories of the wild-type *E. coli* (strain RP437) as a function of time. From a power law fit of the MSD, we estimate that it will take unexposed bacteria at least 30 s to reach the field of view, although the time will vary for different strains depending on the swimming velocity and tumbling frequency. Based on this estimate we only consider the portions of tumble bias and velocity time traces within 30 s after the light has been turned off or on.

#### Run-tumble assignment

To assign run and tumble states we used a Hidden Markov Model with Gaussian-distributed emissions similar to the one described by Dufour *et al.* (46). In this type of model, the state of the system (e.g. ‘run’ or ‘tumble’) is not directly observed (‘hidden’), but its outputs (‘emissions’) such as velocity, acceleration, and angular acceleration can be observed. The hidden states and known outputs are related by an emission probability, while the transition between states is given by a transition matrix.

We implemented the Hidden Markov model using the Python package *hmmlearn* to infer the sequence of ‘hidden’ swimming states from time traces of the observable parameters (47). Parameters of the model—the transition probability matrix and the emission probabilities of the observables—are estimated from a reference dataset consisting of >20,000 pre-stimulus trajectories of wild-type bacteria. Training is done iteratively. At each iteration, velocities and accelerations are normalized by the average swimming velocity (the 95th percentile of the velocity is used over the first iteration), model parameters are estimated from a resulting sequence of observables and an optimal sequence of states is inferred. The process is repeated until the change in normalized velocity between two consecutive iterations is below 2%. To account for the variation in swimming velocity within one trajectory, we used a model with three states—‘fast run’, ‘slow run’, and ‘tumble’. The running velocity was calculated from states corresponding to a fast run. While we performed run-tumble assignment for all the collected trajectories, only those longer than 100 frames were used for all the plots in the paper, as shorter trajectories do not contain enough points for accurate run-tumble assignment as shown by Dufour *et al.* (46).

We validated our assignment by comparing tumble bias time traces obtained with the above analysis to those obtained with alternative run-tumble assignment criteria introduced by Alon *et al*. (48). Under these alternate criteria, bacteria were considered to be tumbling if the velocity was below the 90th percentile of velocities divided by two and if the angular velocity was above 6 rad/s. We compared tumble bias traces obtained with each assignment method. Although the absolute value of tumble bias was offset depending on the specific method used to assign runs and tumbles, the trends—response and adaptation to light exposure—did not depend on the analysis method. Finally, following Wright *et al*. we used angular velocity or RCD (7) (rate of change in direction), which is proportional to tumble bias over a range of values (49), as a population measure of the *E. coli* chemotactic response. We found similar trends in the RCD traces as well. Therefore, we conclude that our results are robust to the method of analysis. The Python library that we developed for detection and analysis of trajectories is available on the GitHub repository: https://github.com/tatyana-perlova/py-taxis.

For each movie, the average tumble bias across all trajectories was calculated in each movie frame. Tumble bias traces from movies taken under the same conditions were averaged, yielding a smooth tumble bias trace as a function of time for each strain or condition. We used standard error between the tumble biases of individual bacterial trajectories as a measure of uncertainty of average tumble bias (Fig. 1C, shaded area). The amplitude *A* was defined as the maximum change in tumble bias upon light exposure compared to the pre-stimulus value and was calculated over a 0.5-s window to minimize the contribution of noise (Fig. 2B). For extracting response and adaptation times, tumble bias response traces during light exposure were fitted to the equation:

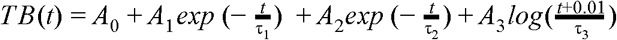

The logarithmic term was added to account for the slow adaptation kinetics in some of the receptor mutants. The response time *t*_*resp*_ was defined as the time it takes for the resulting fitted function to reach 90±1% of the amplitude (Fig. 2B) or:

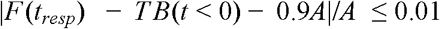

Similarly, the adaptation time *t*_*adap*_ was defined as the time it takes for the difference between the pre-stimulus value of tumble bias and the fitting function to reach 50±1% of the amplitude (Fig. 2B). The error in determining response and adaptation time was calculated as the standard error of the tumble bias at *t*_*resp*_ or *t*_*adap*_ (width of the shaded area around the tumble bias trace, Figure 2) divided by the slope of the linear fit to the trace at that point. For plotting, tumble bias time traces were averaged using a 10-frame moving window.

#### Running velocity analysis

The running velocity was defined as the velocity during frames assigned to runs. For the non-tumbling receptorless strain UU1250 we used velocity during every frame. Time traces of the normalized run velocity (Fig. 4A) were calculated as follows: each individual velocity trace was divided by the pre-stimulus velocity calculated in a 20-s window prior to light exposure. Traces for were then shifted to align ‘light on’ or ‘light off’ frames and averaged using a 10-frame non-overlapping rolling window. As the tumble assignment procedure is not perfect, not all tumbles are detected, which is reflected in a sudden decrease in swimming velocity right after light exposure for strains with a tumbling response to light (data not shown). To measure the effect of light on the running velocity while minimizing the effect of false negatives we use the following procedure: for each individual movie velocities are calculated in a 4-s windows after 30 s of light exposure and then 10, 20 and 30 s after the light is turned off (Figure 4B). Therefore we do not take into account velocity right after the light is turned on. In Aer-only and Trg-only strains, tumble bias does not return to pre-stimulus values even after 30 s of exposure. Therefore these strains were excluded from this analysis. Observation was limited to 30 s to avoid contribution of unexposed bacteria swimming into the field of view. Velocities calculated in this way for each trace were then normalized to the pre-stimulus velocity calculated in a 20-s window and averaged across all the traces for a particular strain. Decrease and recovery of normalized run velocity were calculated as illustrated in Fig. 4B by averaging data points for different strains. The velocity decrease, Δ*v*_*on*_, was an average decrease of normalized run velocity after 30 s of light exposure. The velocity recovery characterizes the fraction of Δ*v*_*on*_ recovered 30 s after light turn-off and was calculated as (Δ*v*_*on*_ - Δ*v*_*off*_)/Δ*v*_*on*_ (Fig. 4B).

## Acknowledgements

We would like to thank Professors Robert Gennis, Christopher Rao, George Ordal and Yann Dufour for advice and discussion. We thank all members of Chemla lab and Gruebele group, and especially Roshni Bano for help and suggestions throughout the course of this work. We thank Anton Goloborodko for help with data analysis. Mutant strains used in this study were kindly provided by Professor John Parkinson. Funding was provided by National Science Foundation grants PHY-1430124 [Physics Frontier Center (PFC) – Center for the Physics of Living Cells (CPLC)] and PHY-1519407 [NSF-Inspire].

## Supplementary materials

Blue light is a universal tactic signal for *Escherichia coli*

Tatyana Perlova, Martin Gruebele and Yann R. Chemla

**Supplementary Figure 1.**
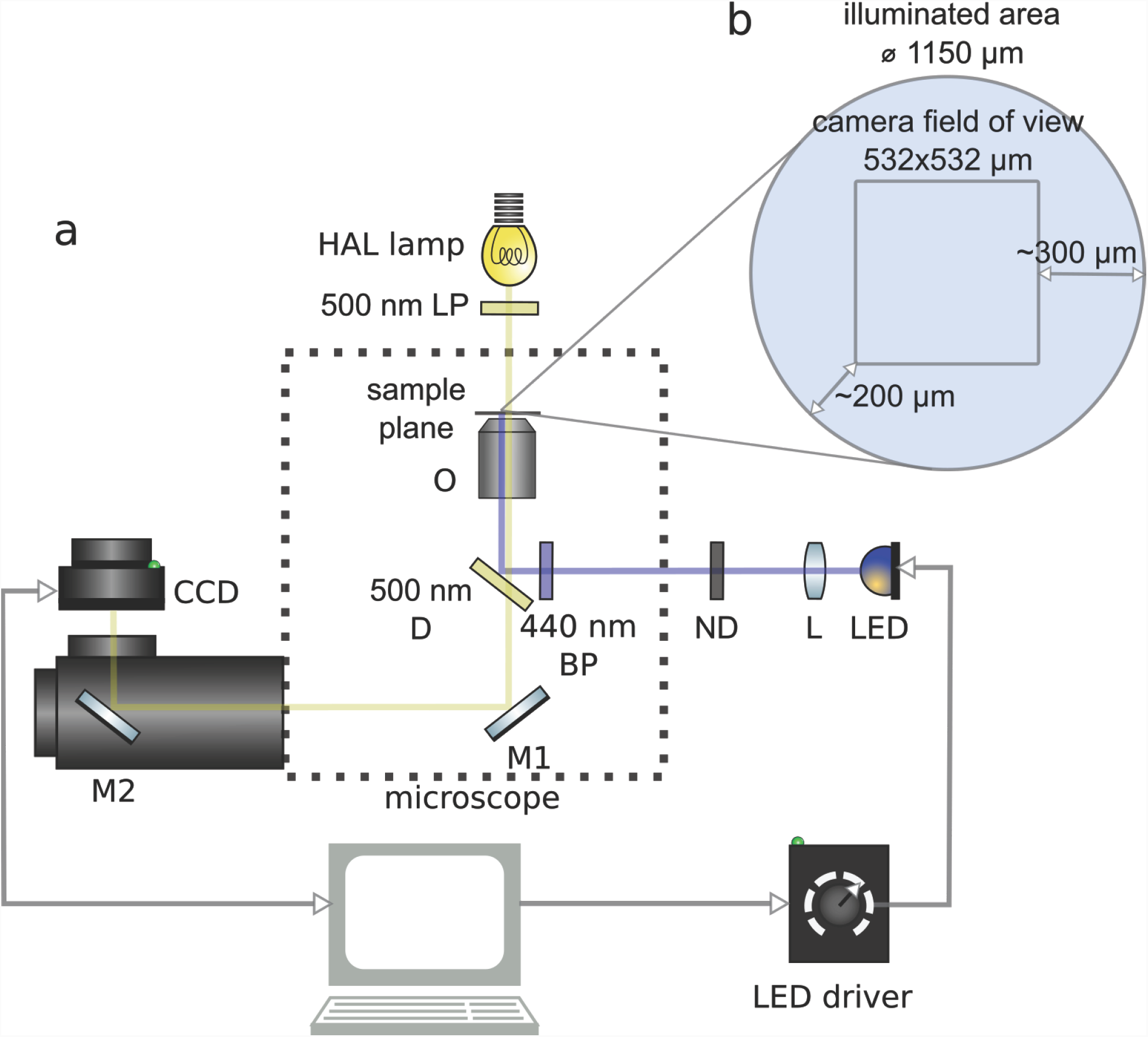
Experimental setup. (A) Schematic of the setup with the light paths indicated by yellow (wide-range visible light from HAL lamp) and blue (blue LED) lines. Components are labeled as follows: LP - long-pass filter, O - 20x objective, D - dichroic, BP - bandpass filter, ND - neutral density filter, L - collimating lens, LED - light emitting diode, M1, M2 - fully reflective mirrors, CCD - Charge-Coupled Device Camera. LED power output is controlled by the current from the LED driver, which is controlled by modulating the voltage through a LabView interface. (B) Area captured by the camera compared to the total illumination area, which is equal to the objective’s field of view. Minimal and maximal distances that unexposed bacteria need to swim to reach the observation area are shown.

**Supplementary Figure 2.**
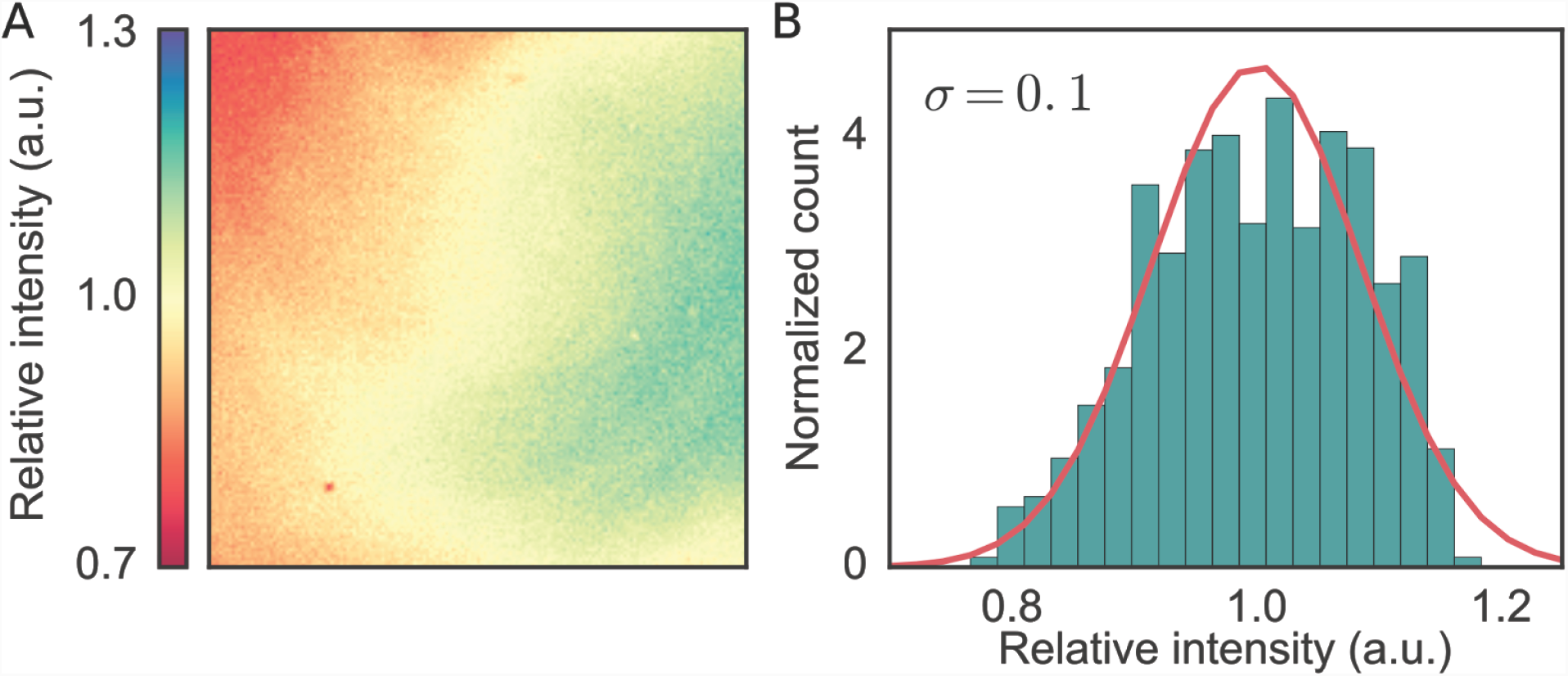
Estimating evenness of the Illumination profile from the brightness of the image captured by the camera. (A) Blue light illumination profile captured by the CCD camera. Color indicates normalized brightness of each pixel (brightness divided by the average brightness). (B) Distribution of the normalized pixel brightness. The standard deviation of the distribution is ∼ 0.1.

**Supplementary Figure 3.**
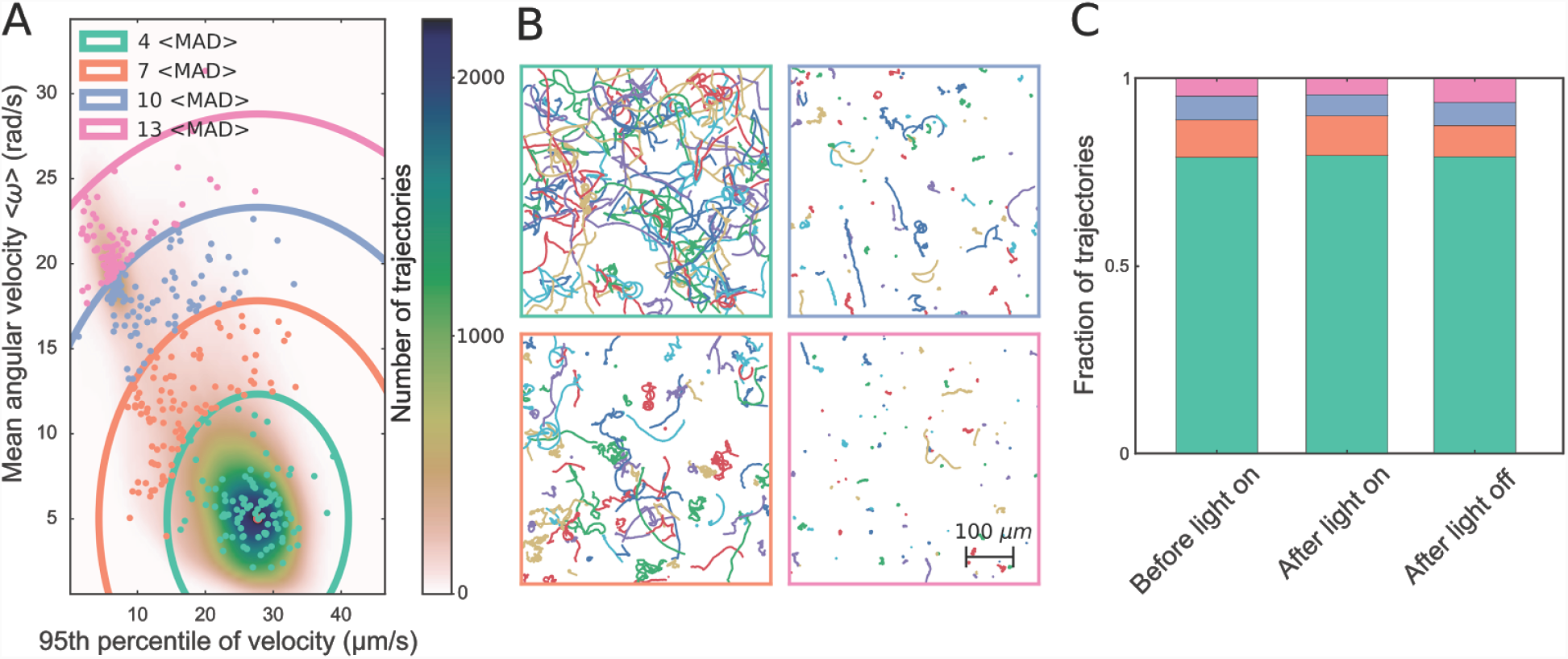
Filtering spurious trajectories. (A) Representative two-dimensional distribution of ∼200,000 trajectories from wild-type *E. coli* plotted against the 95th percentile of velocity and mean angular velocity. Trajectories outside the green contour with radius *R* = 4 <*MAD*> are removed from further analysis (Materials and Methods). Dots indicate randomly selected trajectories shown in panel (B). (B) Each panel contains 100 randomly selected trajectories from within each contour in panel (A). The color of the panel frame indicates which region the trajectories come from in panel (A). (C) Fraction of trajectories within each contour as indicated by the color, before, during, and after light exposure.

**Supplementary Figure 4.**
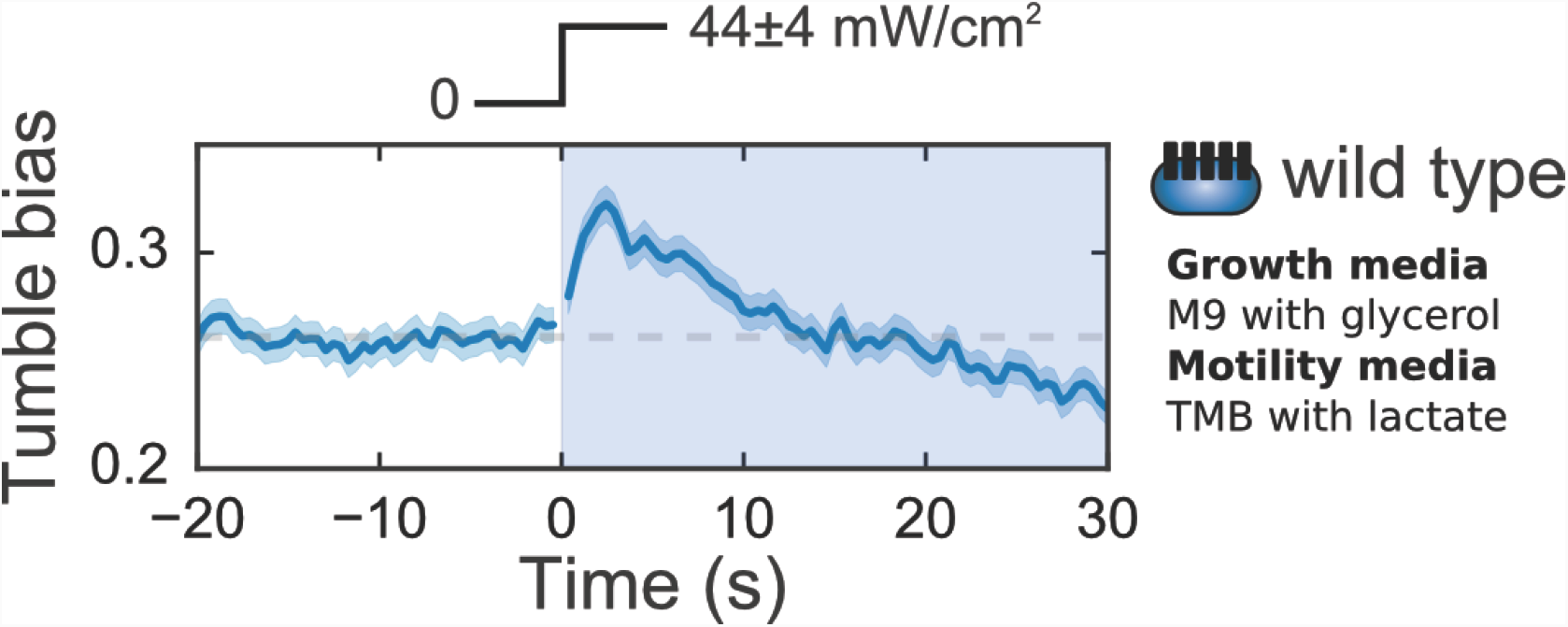
Blue light response under different growth conditions. Tumble bias trace for the wild-type strain RP437 grown in M9 minimal medium with glycerol and resuspended in motility buffer with lactate (Materials and Methods). ∼3000 trajectories were used to calculated average tumble bias at each time point. Response to a turn-on of blue light of intensity 44 ± 4 mW/cm^2^ reproduces previous results on *E. coli* grown with the same carbon source (1).

**Supplementary Figure 5.**
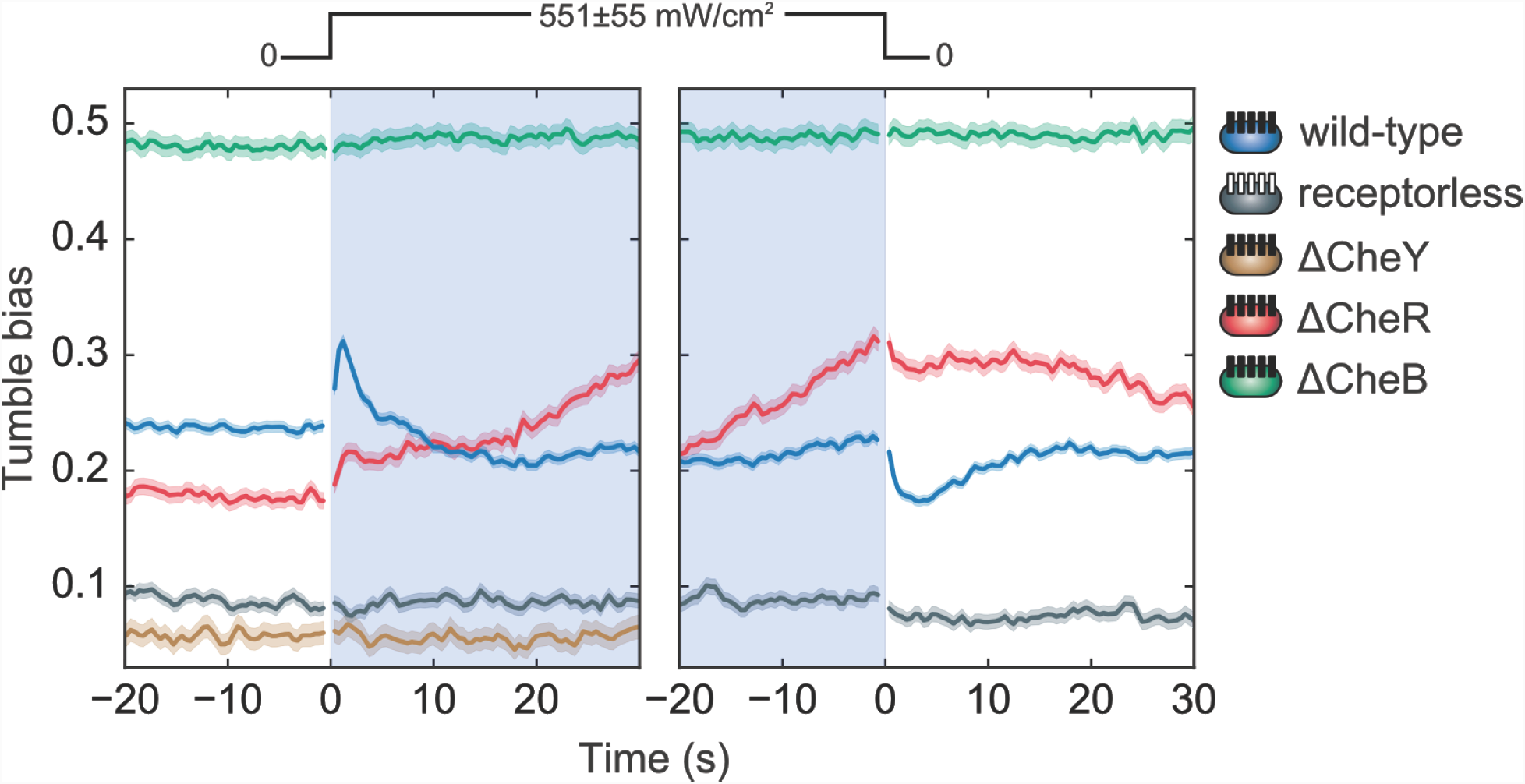
Response to light in *E. coli* is mediated by the chemotaxis network. Tumble bias traces for *E. coli* mutants lacking different components of the chemotaxis network (Table 1): receptorless strain with all receptor types deleted (grey) (2), ΔCheY strain lacking functional CheY (yellow) (3), ΔCheR strain lacking methylesterase CheR (red), ΔCheB strain lacking methyltransferase CheB (green). ∼500 - 3000 trajectories were used to calculated average tumble bias at each time point. The wild-type response trace is shown for comparison (blue). All responses were measured at a blue light intensity of 551 ± 55 mW/cm^2^ as indicated by the light intensity profile.

**Supplementary Figure 6.**
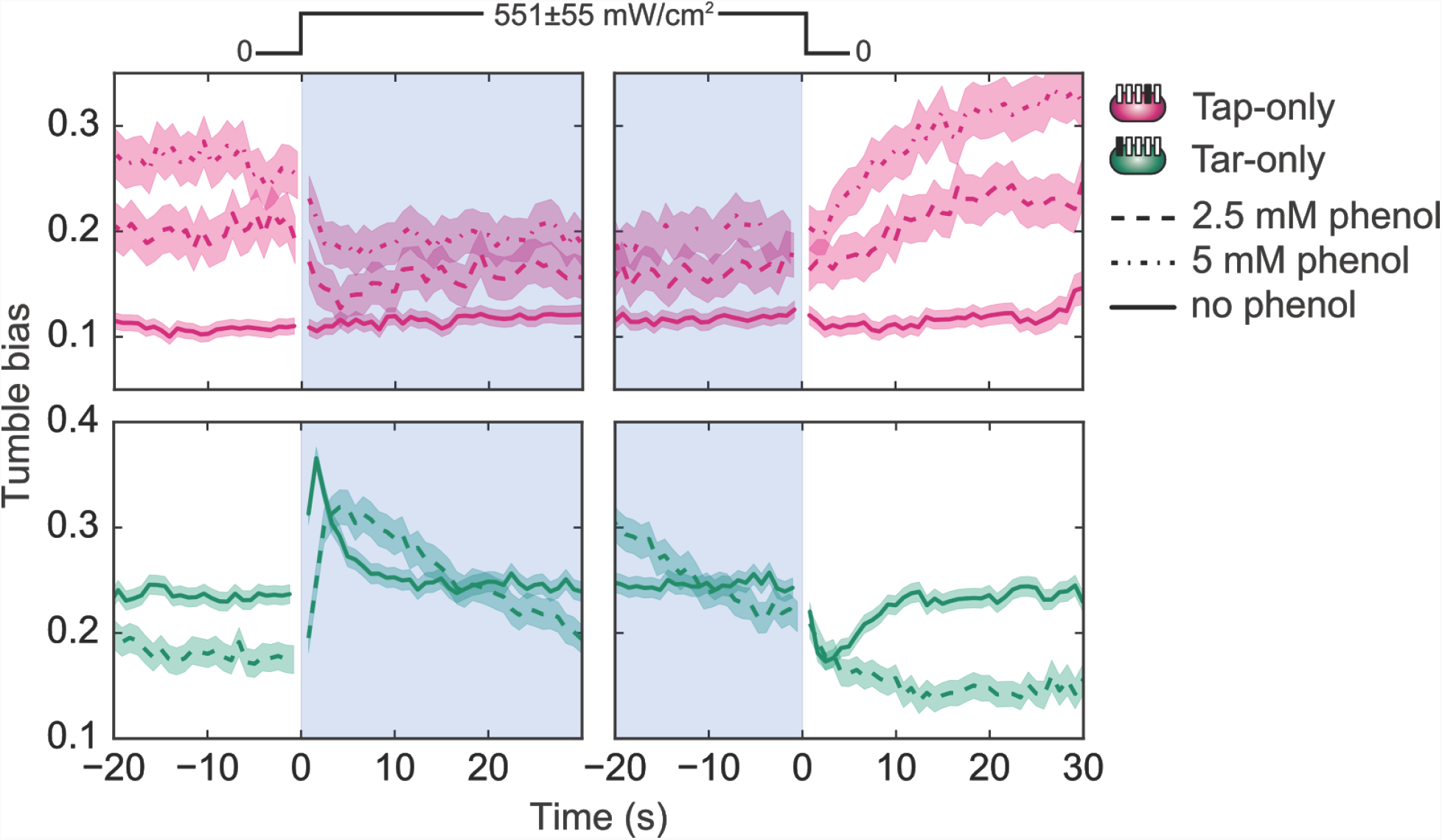
Effect of phenol on the light response in Tap-only and Tar-only strain. Tumble bias traces for Tap-only (magenta) and Tar-only (green) strains without phenol (solid line), with 2.5 mM phenol (dashed line), and with 5 mM phenol (dotted line). ∼200 - 2000 trajectories were used to calculated average tumble bias at each time point.

**Supplementary Figure 7.**
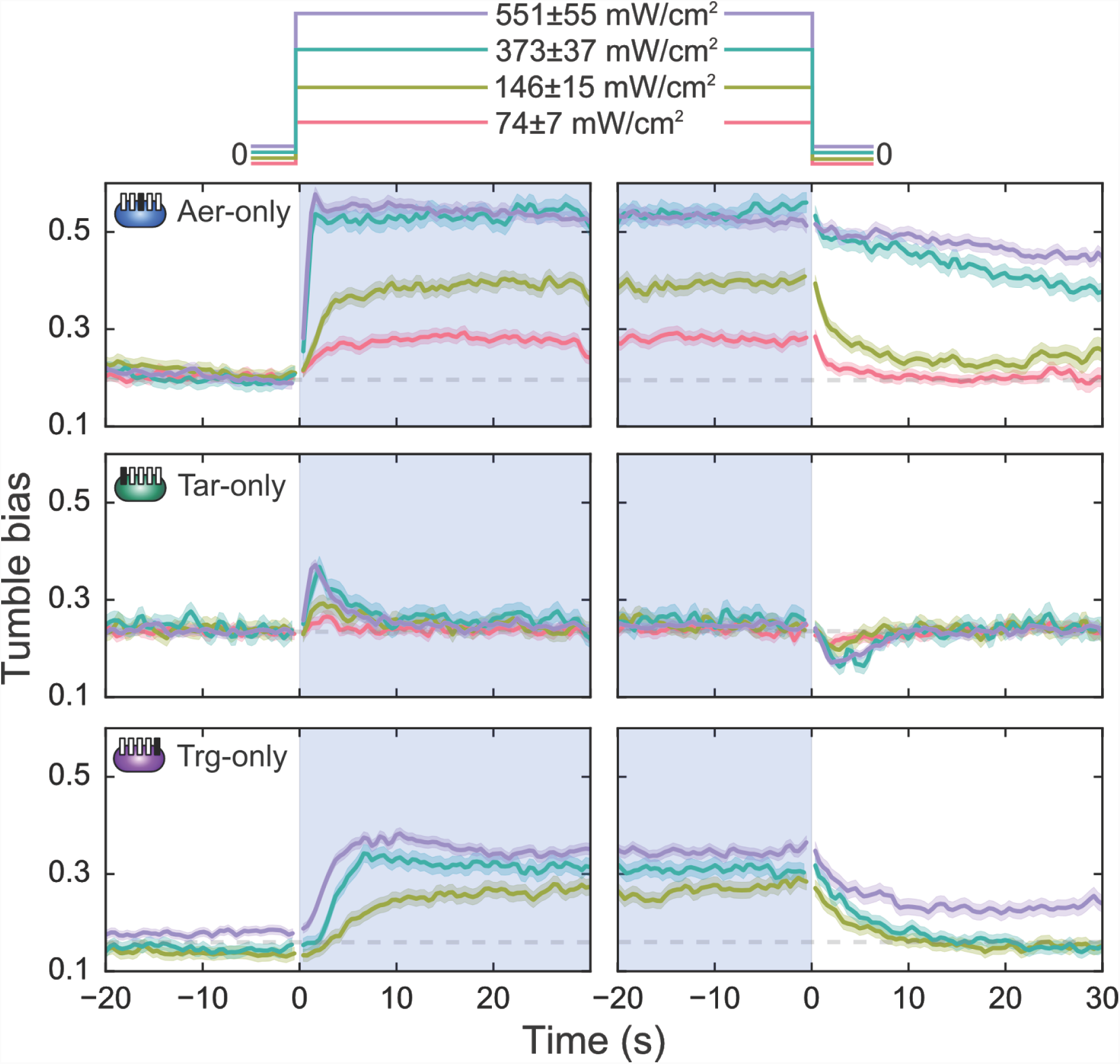
Blue light responses at different intensity levels for Aer-only, Tar-only, and Trg-only strains. Light intensity is indicated by the trace color. Grey dashed lines show prestimulus tumble bias for turn-on and turn-off responses. Light exposure is indicated by the shaded area as well as by the light intensity profile above the plot. ∼200 - 2000 trajectories were used to calculated average tumble bias at each time point.

## References

1. Häder DP. 1987. Photosensory behavior in procaryotes. Microbiol Rev 51:1–21.

2. Armitage JP, Hellingwerf KJ. 2003. Light-induced behavioral responses (‘phototaxis’) in prokaryotes. Photosynth Res 76:145–155.

3. Purcell EB, Crosson S. 2008. Photoregulation in prokaryotes. Curr Opin Microbiol 11:168–178.

4. Taylor BL, Koshland DE. 1975. Intrinsic and extrinsic light responses of Salmonella typhimurium and Escherichia coli. J Bacteriol 123:557–569.

5. Taylor BL, Miller JB, Warrick HM, Koshland DE. 1979. Electron acceptor taxis and blue light effect on bacterial chemotaxis. J Bacteriol 140:567–573.

6. Yang H, Inokuchi H, Adler J. 1995. Phototaxis away from blue light by an Escherichia coli mutant accumulating protoporphyrin IX. Proc Natl Acad Sci U S A 92:7332–7336.

7. Wright S, Walia B, Parkinson JS, Khan S. 2006. Differential activation of Escherichia coli chemoreceptors by blue-light stimuli. J Bacteriol 188:3962–3971.

8. Mears PJ, Koirala S, Rao CV, Golding I, Chemla YR. 2014. Escherichia coli swimming is robust against variations in flagellar number. Elife 1–18.

9. Rebbapragada A, Johnson MS, Harding GP, Zuccarelli AJ, Fletcher HM, Zhulin IB, Taylor BL. 1997. The Aer protein and the serine chemoreceptor Tsr independently sense intracellular energy levels and transduce oxygen, redox, and energy signals for Escherichia coli behavior. Proceedings of the National Academy of Sciences 94:10541–10546.

10. Sourjik V, Wingreen NS. 2012. Responding to chemical gradients: bacterial chemotaxis. Curr Opin Cell Biol 24:262–268.

11. Grebe TW, Stock J. 1998. Bacterial chemotaxis: the five sensors of a bacterium. Curr Biol 8:R154–7.

12. Yang Y, Sourjik V. 2012. Opposite responses by different chemoreceptors set a tunable preference point in Escherichia coli pH taxis. Mol Microbiol 86:1482–1489.

13. Sourjik V, Wingreen NS. 2012. Responding to chemical gradients: bacterial chemotaxis. Curr Opin Cell Biol 24:262–268.

14. Mao H, Cremer PS, Manson MD. 2003. A sensitive, versatile microfluidic assay for bacterial chemotaxis. Proc Natl Acad Sci U S A 100:5449–5454.

15. Paster E, Ryu WS. 2008. The thermal impulse response of Escherichia coli. Proc Natl Acad Sci U S A 105:5373–5377.

16. Taylor BL, Zhulin IB, Johnson MS. 1999. Aerotaxis and other energy-sensing behavior in bacteria. Annu Rev Microbiol 53:103–128.

17. Baker MD, Wolanin PM, Stock JB. 2006. Signal transduction in bacterial chemotaxis. Bioessays 28:9–22.

18. Berg HC. 2000. Motile Behavior of Bacteria. Phys Today 53:24.

19. Hazelbauer GL, Falke JJ, Parkinson JS. 2008. Bacterial chemoreceptors: high-performance signaling in networked arrays. Trends Biochem Sci 33:9–19.

20. Vladimirov N, Sourjik V. 2009. Chemotaxis: how bacteria use memory. Biol Chem 390:1097–1104.

21. Li M, Hazelbauer GL. 2004. Cellular Stoichiometry of the Components of the Chemotaxis Signaling Complex 186:3687–3694.

22. Barnakov AN, Barnakova LA, Hazelbauer GL. 1998. Comparison in vitro of a high- and a low-abundance chemoreceptor of Escherichia coli: Similar kinase activation but different methyl-accepting activities. J Bacteriol 180:6713–6718.

23. Yamamoto K, Macnab RM, Imae Y. 1990. Repellent response functions of the Trg and Tap chemoreceptors of Escherichia coli. J Bacteriol 172:383–388.

24. Berman - Academic Press IB, York N, 1971. 1971. Handbook of fluorescence spectra of aromatic molecules.

25. Yuan J, Branch RW, Hosu BG, Berg HC. 2012. Adaptation at the output of the chemotaxis signalling pathway. Nature 484:233–236.

26. Bibikov SISI, Miller ACAC, Gosink KK, Parkinson JS. 2004. Methylation-independent aerotaxis mediated by the Escherichia coli Aer protein. J Bacteriol 186:3730–3737.

27. Vladimirov N, Løvdok L, Lebiedz D, Sourjik V. 2008. Dependence of bacterial chemotaxis on gradient shape and adaptation rate. PloS Comput Biol 4:e1000242.

28. Min TL, Mears PJ, Golding I, Chemla YR. 2012. Chemotactic adaptation kinetics of individual Escherichia coli cells. Proceedings of the National Academy of Sciences 109:1–6.

29. Parkinson JS. 2004. Crosslinking snapshots of bacterial chemoreceptor squads 101:6314–6318.

30. Briegel A, Li X, Bilwes AM, Hughes KT, Jensen GJ, Crane BR. 2012. Bacterial chemoreceptor arrays are hexagonally packed trimers of receptor dimers networked by rings of kinase and coupling proteins. Proceedings of the National Academy of Sciences 109:3766–3771.

31. Parkinson JS, Hazelbauer GL, Falke JJ. 2015. Signaling and sensory adaptation in Escherichia coli chemoreceptors: 2015 update. Trends Microbiol 1–10.

32. Sourjik V, Berg HC. 2002. Receptor sensitivity in bacterial chemotaxis. Proc Natl Acad Sci U S A 99:123–127.

33. Taylor BL. 2007. Aer on the inside looking out: paradigm for a PAS-HAMP role in sensing oxygen, redox and energy. Mol Microbiol 65:1415–1424.

34. White David. 2012. Membrane Bioenergetics: The Proton Potential. PloS Comput Biol 10:e1003870.

35. Fung DC, Berg HC. 1995. Powering the flagellar motor of Escherichia coli with an external voltage source. Nature.

36. Gabel CV, Berg HC. 2003. The speed of the flagellar rotary motor of Escherichia coli varies linearly with protonmotive force. Proc Natl Acad Sci U S A 100:8748–8751.

37. Nakamura S, Nakamura T, Ogura Y. 1963. Absorption spectrum of flavin mononucleotide semiquinone. J Biochem 53:143–146.

38. Losi A, Gärtner W. 2011. Old chromophores, new photoactivation paradigms, trendy applications: flavins in blue light-sensing photoreceptors. Photochem Photobiol 87:491–510.

39. Conrad KS, Manahan CC, Crane BR. 2014. Photochemistry of flavoprotein light sensors. Nat Chem Biol 10:801–809.

40. Kao Y-T, Saxena C, He T-F, Guo L, Wang L, Sancar A, Zhong D. 2008. Ultrafast dynamics of flavins in five redox states. J Am Chem Soc 130:13132–13139.

41. Edwards JC, Johnson MS, Taylor BL. 2006. Differentiation between electron transport sensing and proton motive force sensing by the Aer and Tsr receptors for aerotaxis. Mol Microbiol 62:823–837.

42. Blount ZD. 2015. The unexhausted potential of E. coli. Elife 4.

43. Kalinin Y, Neumann S, Sourjik V, Wu M. 2010. Responses of Escherichia coli bacteria to two opposing chemoattractant gradients depend on the chemoreceptor ratio. J Bacteriol 192:1796–1800.

44. Bradski G. 2000. Open Source Computer Vision Library. Dr Dobb’s Journal of Software Tools.

45. Allan D, Caswell TA, Keim N, Boulogne F, Perry RW, Uieda L. 2014. trackpy: Trackpy v0.3.0.

46. Dufour YS, Gillet S, Frankel NW, Weibel DB, Emonet T. 2016. Direct Correlation between Motile Behavior and Protein Abundance in Single Cells. PloS Comput Biol 12:e1005041.

47. Pedregosa F, Varoquaux G, Gramfort A, Michel V, Thirion B, Grisel O, Blondel M, Prettenhofer P, Weiss R, Dubourg V, Vanderplas J, Passos A, Cournapeau D, Brucher M, Perrot M, Duchesnay É. 2011. Scikit-learn: Machine Learning in Python. J Mach Learn Res 12:2825–2830.

48. Alon U, Camarena L, Surette MG, Aguera y Arcas B, Liu Y, Leibler S, Stock JB. 1998. Response regulator output in bacterial chemotaxis. EMBO J 17:4238–4248.

49. Khan S, Castellano F, Spudich JL, McCray JA, Goody RS, Reid GP, Trentham DR. 1993. Excitatory signaling in bacterial probed by caged chemoeffectors. Biophys J 65:2368–2382.

50. Parkinson JS, Houts SE. 1982. Isolation and Behavior of Escherichia coli Deletion Mutants Lacking Chemotaxis Functions.

51. Min TL, Mears PJ, Chubiz LM, Rao CV, Golding I, Chemla YR. 2009. High-resolution, long-term characterization of bacterial motility using optical tweezers. Nat Methods 6:831–835.

52. Gosink KK, Burón-Barral M del C, Parkinson JS. 2006. Signaling interactions between the aerotaxis transducer Aer and heterologous chemoreceptors in Escherichia coli. J Bacteriol 188:3487–3493.

53. Bibikov SI, Biran R, Rudd KE, Parkinson JS. 1997. A signal transducer for aerotaxis in Escherichia coli. J Bacteriol 179:4075–4079.

54. Wright S, Walia B, Parkinson JS, Khan S. 2006. Differential activation of Escherichia coli chemoreceptors by blue-light stimuli. J Bacteriol 188:3962–3971.

55. Bibikov SI, Miller AC, Gosink K, Parkinson JS. 2004. Methylation-independent aerotaxis mediated by the Escherichia coli Aer protein. J Bacteriol 186:3730–3737.

56. Min TL, Mears PJ, Chubiz LM, Rao CV, Golding I, Chemla YR. 2009. High-resolution, long-term characterization of bacterial motility using optical tweezers. Nat Methods 6:831–835.

